# Zebrafish Galectin 3 binding protein (Lgals3bp) is the target antigen of the microglial 4C4 monoclonal antibody

**DOI:** 10.1101/2022.04.05.486973

**Authors:** Mireia Rovira, Alice Montanari, Latifa Hammou, Laura Chomette, Magali Miserocchi, Jennifer Pozo, Virginie Imbault, Xavier Bisteau, Valérie Wittamer

## Abstract

**Background:** Two decades ago, the fish-specific monoclonal antibody 4C4 was found to be highly reactive to zebrafish microglia, the macrophages of the central nervous system. This has resulted in 4C4 being widely used, in combination with available fluorescent transgenic reporters to identify and isolate microglia. However, the target protein of 4C4 remains unidentified, which represents a major caveat. In addition, whether the 4C4 expression pattern is strictly restricted to microglial cells in zebrafish has never been investigated.

**Results:** Having demonstrated that 4C4 is able to capture its native antigen from adult brain lysates, we used immunoprecipitation/mass-spectrometry, coupled to recombinant expression analyses, to identify its target. The cognate antigen was found to be a paralog of Galectin 3 binding protein (Lgals3bpb), known as MAC2-binding protein in mammals. Notably, 4C4 did not recognize other paralogs, demonstrating specificity. Moreover, our data show that Lgals3bpb expression, while ubiquitous in microglia, also identifies leukocytes in the periphery, including populations of gut and liver macrophages.

**Conclusions:** The 4C4 monoclonal antibody recognizes Lgals3bpb, a predicted highly glycosylated protein whose function in the microglial lineage is currently unknown. Identification of Lgals3bpb as a new pan-microglia marker will be fundamental in forthcoming studies using the zebrafish model.

## INTRODUCTION

Microglia are the resident macrophages in the central nervous system (CNS). They play key functions in maintaining brain homeostasis through their constant surveillance of the brain parenchyma and their interactions with other cells in the CNS. Microglia have been shown to exhibit both neuroprotective and neurotoxic functions and have important roles in brain diseases such as amyotrophic lateral sclerosis, multiple sclerosis, Parkinson’s disease, Alzheimer’s disease, glioma, or HIV-related dementia ^1,2^. While our current understanding of microglia biology is mainly derived from investigations performed in mice, the zebrafish has also emerged over the past few years as a powerful complementary model to study microglia ^3–5^. Studies in zebrafish have provided new clues on the cellular and molecular requirements for microglia ontogeny and cell functions and have contributed to shedding light on evolutionary aspects of microglia properties across vertebrate species.

In zebrafish, the study of microglia largely relies on the use of a variety of fluorescent reporter lines allowing to take full advantage of the transparency of the embryo to directly visualize microglial cells in their microenvironment. Antibodies labelling microglia in zebrafish have been limited since they are raised against mammalian proteins and usually display low cross-reactivity with zebrafish proteins. Among these, the 7.4.C4 monoclonal antibody, available as an hybridoma source (ECACC 92092321, deposited by A. Dowding from the King’s College London, UK) and commonly referred to as “4C4”, has been extensively used as a “fish macrophage/microglia-specific” antibody ^6–8^. According to the ECACC datasheet, 4C4 was originally raised against protein extracts derived from optic nerves of the freshwater fish *Oreochromis mossambicus* at 12 days post injury. It was first used in zebrafish about 20 years ago, and suggested in immunofluorescence experiments to be highly specific for microglia, based on morphological criteria ^6,9^. Since then, a growing number of publications have promoted 4C4 as an invaluable tool for the study of zebrafish microglia, and its use has been extended to flow cytometry as well as labeling of macrophages outside the CNS ^10^. Despite its wide use, the 4C4 antibody has remained poorly characterized and, more importantly, its target protein is unknown. In this study, we sought to overcome these limitations. We report the identification of Galectin 3 binding protein (Lgals3bp) as the 4C4 antigen, using proteomic, molecular and genetic approaches. We further show that Lgals3bp is specifically expressed in microglia within the brain parenchyma across the zebrafish lifespan, and we uncovered 4C4 labelling in discrete populations of tissue macrophages.

## RESULTS

### The 4C4 target antigen is ubiquitously expressed in zebrafish microglia

We first examined the expression profile of the antigenic protein recognized by the 4C4 antibody in microglia across the life span. In the zebrafish embryo, microglia can be easily identified in the developing brain parenchyma based on *mpeg1*:GFP transgene expression ^11,12^. We performed immunostaining for GFP and 4C4 on Tg(*mpeg1*:GFP) embryos at 3 days post fertilization (dpf), a stage where the phenotypic transition to differentiated microglia is completed ^11^. As shown in Figure 1A, all GFP^+^ microglia display 4C4 immunostaining. These observations are consistent with previous reports and suggest that the 4C4 antigen is constitutively and ubiquitously expressed among embryonic microglial cells. Next, we evaluated 4C4 expression in adult microglia. We used the *p2ry12:p2ry12*-GFP line, where a P2ry12-GFP fusion protein is expressed under the control of the *p2ry12* promoter, a well-known canonical microglia marker in mammals and zebrafish ^13,14^. We found that all GFP^+^ cells co-localize with the 4C4 staining, as unveiled by immunofluorescence performed on adult brain sections (Figure 1B). Importantly, staining with the pan-leukocytic marker L-plastin (Lcp1) further confirmed the hematopoietic identity of the 4C4^+^ cells within the adult brain parenchyma. Collectively, these results indicate that the 4C4 antibody labels all zebrafish microglia throughout life, regardless of their developmental origin ^11^.

**Figure 1.**
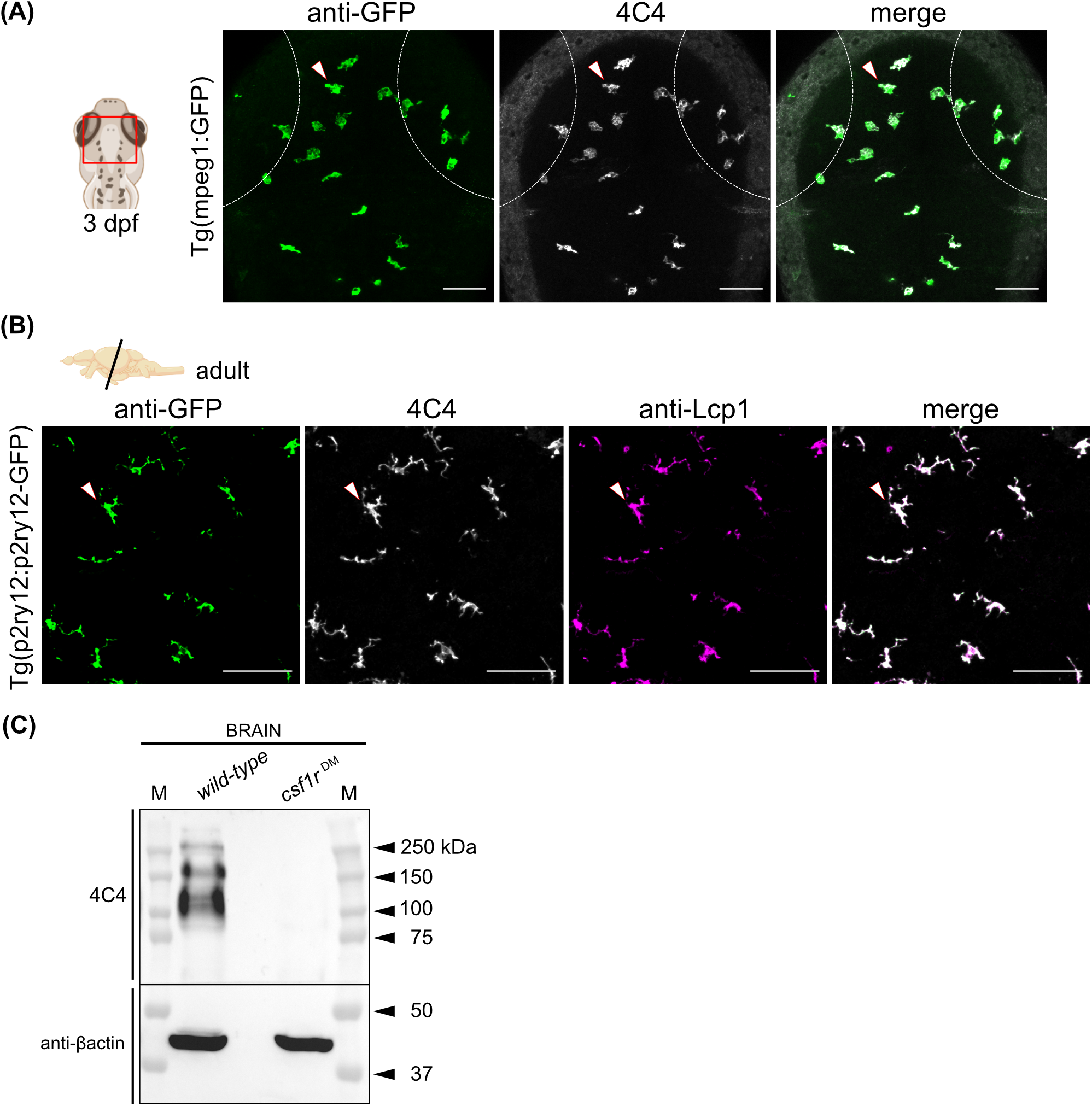
The 4C4 antibody labels microglial cells in the embryonic and adult zebrafish brain. **(A)** Dorsal view (red rectangle) of the optic tectum of a Tg(*mpeg1*:GFP) embryo at 3 dpf co-immunostained with the 4C4 antibody. Anti-GFP (green), 4C4 (grey) and merge of the two channels are shown. Dashed lines represent the eye edges. Images were taken using a 25X water-immersion objective. **(B)** Immunofluorescence on transversal brain sections (14 μm) from adult *Tg(p2ry12-p2ry12:GFP)* zebrafish co-immunostained with 4C4 and Lcp1 (L-plastin) antibodies. Anti-GFP (green), 4C4 (grey), anti-Lcp1 (magenta) and merge of the three channels. Images were taken using a 20X objective. Images in (A) and (B) correspond to orthogonal projections and the white arrowheads point to microglial cells. Scale bars 50 μm. dpf, days post-fertilization. **(C)** Detection of the 4C4 target protein by western blot. Protein lysate from a *wild-type* and a *csf1r*^*DM*^ mutant adult brain, pactin was used as a loading control. M, protein marker.

### The 4C4 antibody recognizes one or several proteins with high molecular weights

As we ultimately sought to isolate and characterize the protein recognized by 4C4, we assessed whether this antibody is suitable for antigen detection in complex mixtures and we tested its performance in immunoblot applications. We prepared protein extracts from the zebrafish adult brain that were separated on a gel under denaturing conditions (SDS-PAGE) and followed by immunoblotting. This revealed several immunoreactive bands at molecular weights between 100-250 kDa (Figure 1C). While it was not possible to determine whether these bands represent variants of the same target protein or different proteins bearing the same epitope, we found that all were notably absent in brain lysates prepared from microglia-deficient *csf1r*^*DM*^ fish (Figure 1C). This is consistent with the microglia-specific expression pattern observed for 4C4 by immunofluorescence in adult brain tissue sections. Collectively, these results indicate the functionality of 4C4 for immunoblot assays, allowing to unveil the existence of one or several microglial target proteins with high apparent molecular weight(s).

### Immunoprecipitation coupled to mass-spectrometry identifies putative 4C4 targets

Our findings that 4C4 binds to its antigen under denaturing conditions indicate this antibody recognizes a linear epitope. We then tested whether 4C4 could also be efficient in precipitating its target. Therefore, to identify the antigen recognized by 4C4, we combined immunoprecipitation (IP) with mass spectrometry (MS) (Figure 2A). Brain protein extracts were incubated with either 4C4 or an isotype-matched control antibody (IgG1_k_) followed by the tryptic digestion of the precipitates. The resulting mixtures of tryptic peptides were then analyzed by liquid chromatography tandem mass spectroscopy (LC-MS/MS). Importantly, western-blotting analysis of the immunoprecipitates prior to enzymatic digestion showed that 4C4 specifically captured proteins in a similar pattern to that seen in direct immunoblotting, with molecular weights between 100-250 kDa. This suggests that the 4C4 antibody displays appreciable immunoprecipitation properties.

**Figure 2.**
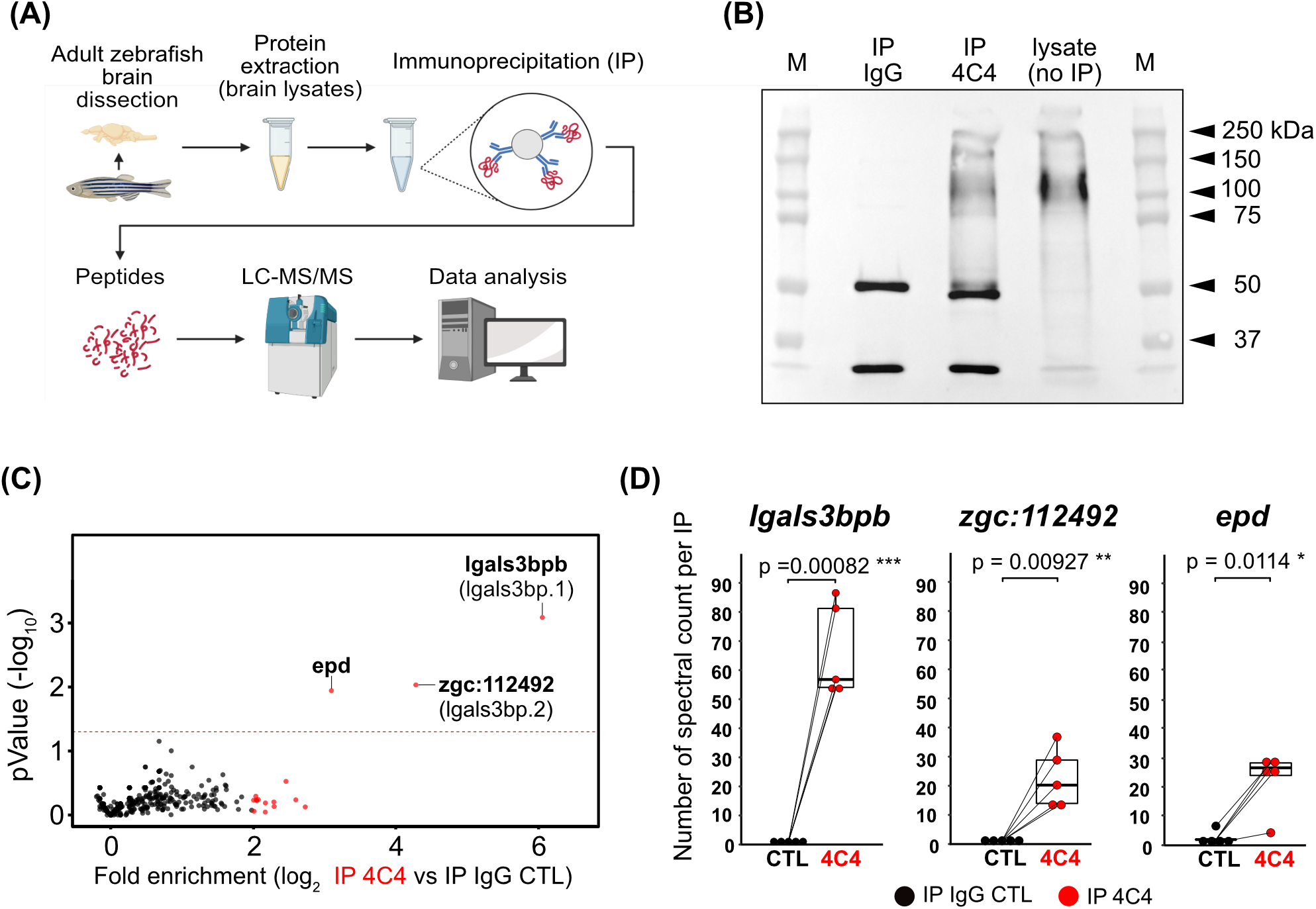
Identification of candidate 4C4 antigens by proteomic analysis. **(A)** Antigen identification strategy, from sample preparation to LC-MS analysis. (B) Immunoprecipitation (IP) of the target protein detected by western blot using the 4C4 antibody. Full blot showing: IP using the control isotype antibody lgG_k1_ (IP IgG), IP using the 4C4 antibody (IP 4C4), brain lysate as a positive control (lysate). The two bands detected in the IP samples correspond to the heavy and light chains (50 and 25 kDa approximately) of the primary antibody that are being recognized by the secondary antibody. M, protein marker; kDa, kilodaltons. **(C)** Volcano plot of averaged enrichment of protein in 4C4 (IP 4C4) versus lgG_k_ control (IP IgG CTL) immunoprecipitations. Red dots: enriched proteins (Fold change ≥ 2) p-val<0.05. (D) Reproducibility boxplot of the total number of spectral counts quantified in each IP linked by pair (n=5 independent experiments) for the 3 most significantly enriched proteins are shown. ***p<0.001, **p<0.01, *p<0.05; paired t-test.

For LC-MS/MS analyses, five independent replicates of each IP experiment were performed and measured per condition. The proteomic analysis identified three proteins that were significantly enriched in the 4C4 samples (Figure 2C, Supplementary Table S1 (http://gofile.me/5Ijcf/DdSQ5Dr8X)). With a log_2_ fold enrichment of 6.05 (p-val=0.0008) and 4.28 (p-val=0.009), respectively, the two most enriched candidates were the products of *galectin 3 binding protein b (lgals3bpb)* and *zgc:112492*, two paralogous genes predicted to encode proteins with a molecular weight of approximately 65 kDa. The third protein of interest was *Ependymin*, a 24 kDa protein showing a log_2_ fold enrichment of 3.10 (p-val=0.01) and less than 25% shared identity with the two other candidates. To show the reproducibility of the different experiments, Figure 2D displays the number of spectral counts obtained from each IP for each of the three significantly identified enriched proteins identified. For every replicate, *lgals3bpb* protein appeared to be more enriched as presented by the higher number of spectral counts than the other two candidates (Figure 2D). This suggests a greater specificity of the 4C4 antibody for *lgals3bpb* than for the other two proteins. Importantly, although the predicted products of *lgals3bpb* and *zgc:112492* share around 80% of identity at the protein level (Figure 3A), we identified eighteen peptides that mapped exclusively with Lgals3bpb, covering 53% of the protein sequence (Figure 3B). Seven additional peptides mapping to conserved regions between the *lgals3bpb* and *zgc:112492* paralogs were also found. Remarkably, however, we only detected a single peptide corresponding to the gene product of *zgc:112492*. For *Ependymin*, we identified five peptides representing 50.7% total coverage. When browsing published datasets for expression ^8,15,16^, we found that amongst the three candidates, only *lgals3bpb* was expressed at high level in microglia. Collectively, our IP-MS approach identified a limited number of proteins of interest. Among these, *Lgals3bpb* appears as a strong candidate for the 4C4 antigen based on the spectral counts, the number of unique peptides, protein sequence coverage, and publicly available expression data.

**Figure 3.**
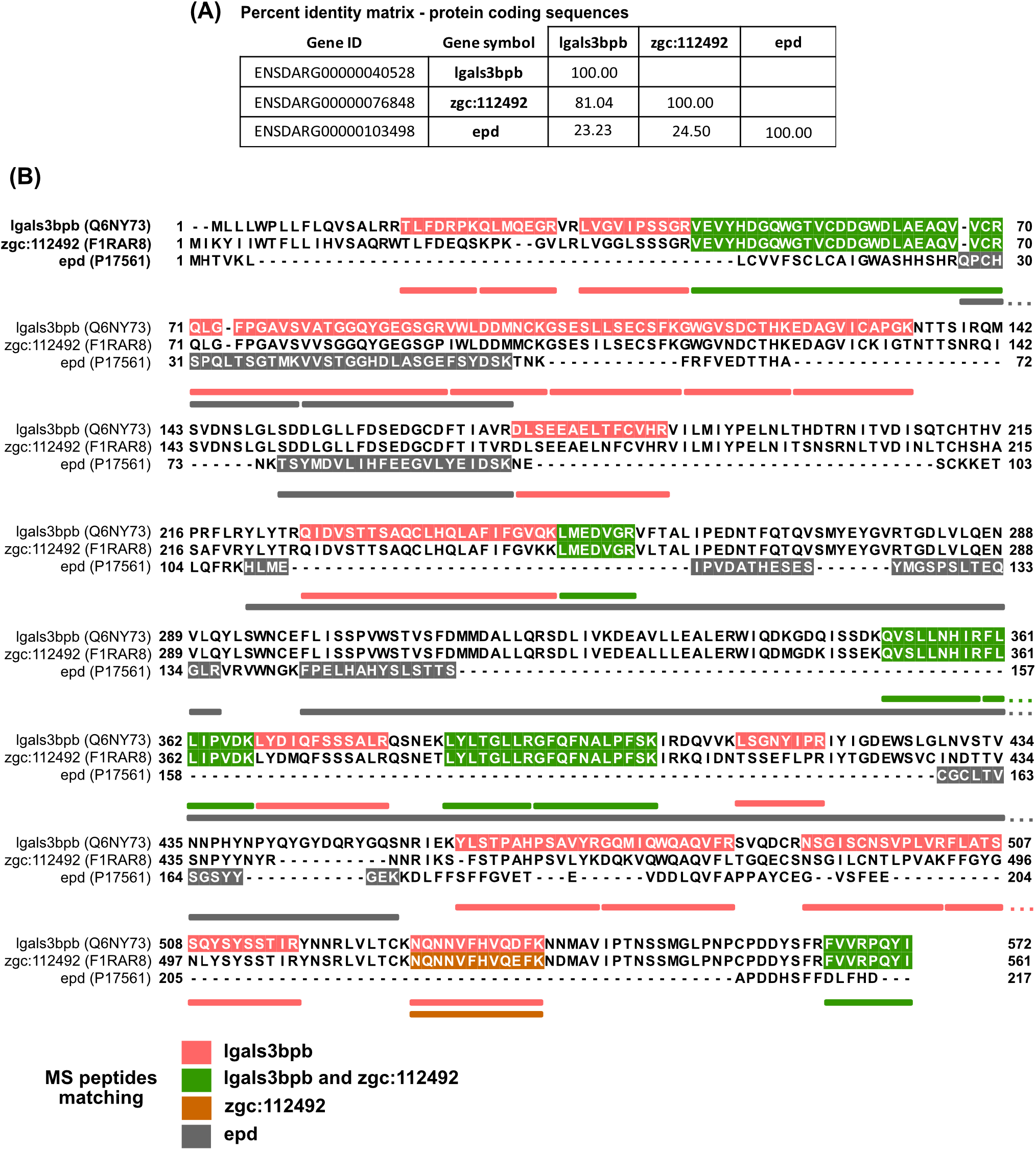
Mapping of the proteotypic peptides shows specificity for the Lgals3bpb sequence. **(A)** Sequence identity (%) between the coding sequences of *Igals3bp, zgc:* 112492 and *epd*. Alignment performed using Clustal Omega (EMBL-EBI) **(B)** Multiple sequence alignment of the coding sequences of *Igals3bpb, zgc*: 112492 and epd with the identified peptides remapped on each sequence. Colors indicate the proteotypicity or group specificity of each identified tryptic peptide. The gene symbol and the UniProtKB identifier are shown.

### The 4C4 antibody recognizes the protein encoded by the *lgals3bpb* gene

To test our hypothesis, we cloned the coding sequence of zebrafish *lgals3bpb*, transiently expressed it in mammalian HEK-293T cells and assessed 4C4 antibody recognition by immunofluorescence. Controls included non-transfected cells and cells transfected with the empty pcDNA3 expression vector. As expected, no signal was detected in controls (Figure 4A-B). In contrast, cells transfected with *lgals3bpb* showed strong heterogenous staining with 4C4, indicating that the antibody detects the overexpressed Lgals3bpb in *lgals3bpb*-expressing cells (Figure 4C-D). Based on these findings, we sought to test whether the gene product of *zgc:112492*, the paralog of *lgals3bpb* also identified in our mass spectrometry analyses, was recognized by 4C4. However, overexpression of the coding sequence of *zgc:112492* in HEK-293T cells did not result in any staining (Figure 4E). These observations prompted us to also assess 4C4 binding to cells engineered to express *lgals3bpa*, a third paralog which was not found in our proteomic analyses but shares 79.5 % identity with *lgals3bpb*. Like for *zgc:112492*, no 4C4 positive signal was observed for this paralog (data not shown). Taken together, these results indicate that *Lgals3bpb*, and not its paralogs, is the target of the 4C4 antibody.

**Figure 4.**
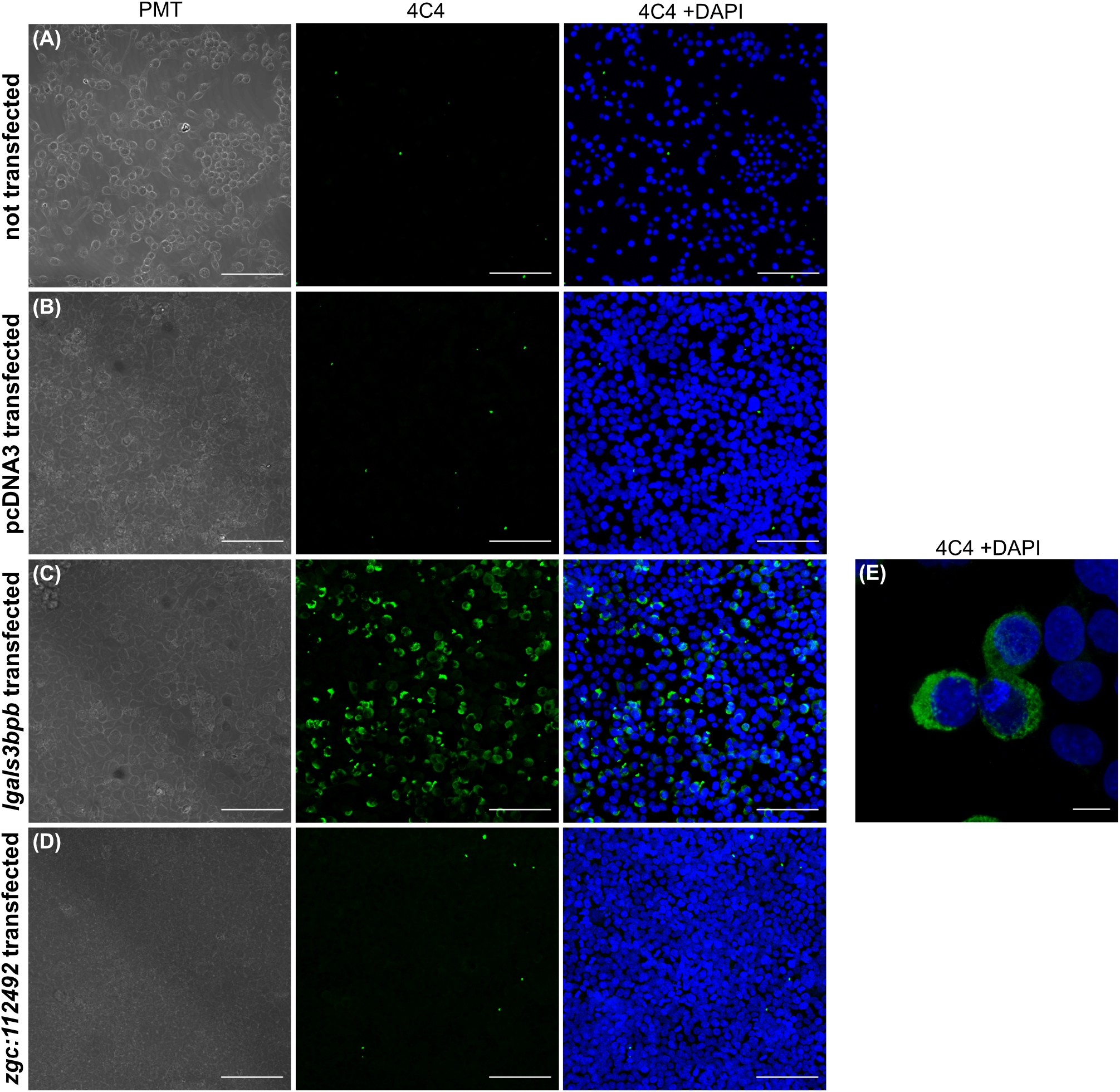
Validation of *Igals3bpb* as the 4C4 antigen by recombinant protein expression in HEK293T cells. **(A-E)** Immunofluorescence of HEK293T cells (A) or following transfection with the empty pcDNA3 plasmid (B) or *Igals3bpb* (C,E) and *zgc:112492* (D) coding sequences. Bright field (left panel), 4C4 staining (middle panel) and 4C4 with DAPI staining (right panel) channels are shown (3 independent experiments). Scale bars 100 μm. Images were taken using a 20X objective. (E) High magnification from pcDNA3-/ga/s3bpb transfected cells. Scale bar 10 μm. Image was taken using a 40X water-immersion objective and a numerical zoom of 3.

### Staining of the 4C4 antibody in peripheral tissues

Having identified Lgals3bpb as the antigen for 4C4, we wondered whether the 4C4 antibody target protein is expressed in other zebrafish tissues in addition to the brain. Previous studies reported that LGALS3BP, the mammalian ortholog of zebrafish *lgals3bpb*, is expressed in different tissues at the mRNA and protein levels. We first investigated the presence of the protein by Western blotting in tissue extracts obtained from various adult zebrafish organs, including brain, eye, liver, intestine, heart, muscle, and whole kidney marrow, the site of hematopoiesis in the adult zebrafish. Consistent with the expression of Lgals3bpb in microglial cells, the 4C4 staining was especially intense in the brain and in the eye (Figure 5A). In other organs, however, the signal was absent or barely detectable. To complement these analyses, we also performed immunostaining with 4C4 on tissue sections from peripheral organs known to contain large numbers of macrophages, such as liver ^17^ and intestine ^18,19^. Samples were prepared from zebrafish carrying the *mpeg1*:GFP transgene, allowing to use GFP as a readout for the presence of macrophages in these tissues, and co-stained with 4C4, anti-GFP and anti-pan-leukocytic Lcp1 antibodies. In the liver, we occasionally found some 4C4^+^ cells, which were all GFP^+^ Lcp1^+^. Intriguingly, these cells were frequently seen in the vicinity of blood vessels, identified on the sections by the presence of erythrocyte nuclei stained with DAPI (Figure 5B-C). Although the *mpeg1* transgene also labels a subpopulation of B cells, we previously showed these cells are scarce in the liver ^18^, indicating 4C4^+^ GFP^+^ cells likely represent macrophages. In the intestine, a proportion of GFP^+^ Lcp1^+^ cells also showed 4C4 immunoreactivity. However, we also identified GFP^-^ Lcp1^+^ 4C4^+^ cells (Figure 5D), suggesting that in this organ, leukocytic Lgasl3bpb expression may not be restricted to *mpeg1*:GFP^+^ immune cells. It should also be noticed that in both organs, not all GFP^+^ cells were labeled with the 4C4 antibody, suggesting that, unlike in microglia, Lgasl3bpb is not ubiquitously expressed in peripheral macrophages.

**Figure 5.**
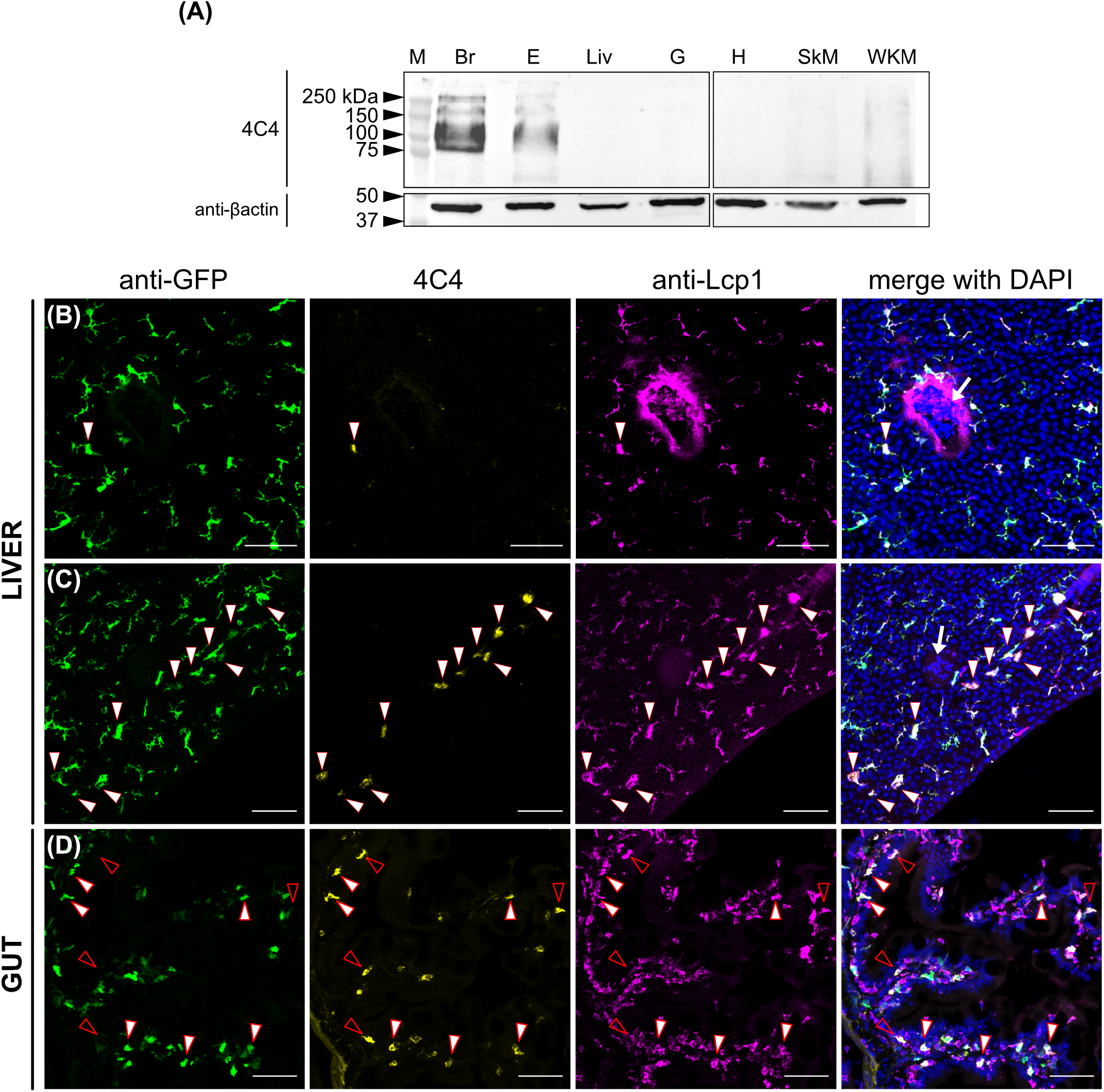
Expression of *Igals3bpb* among adult tissues. **(A)** Tissue distribution by western blot, using the 4C4 antibody and βactin as a loading control. Br, brain; E, eyes; Liv, liver, G, gut; H, heart; SkM, skeletal muscle; WKM, whole kidney marrow (n=3 fish/tissue). kDa, kilodaltons. **(B-D)** Immunofluorescence on liver (C-D) and gut (E) sections (14 µm) from an adult Tg(*mpeg1*:GFP) fish. Anti-GFP (green), 4C4 (yellow), anti- Lcp1 (magenta) and a merge including DAPI staining of the three channels are shown. White arrowheads point to GFP^+^ 4C4^+^ Lcp1^+^ cells while empty arrowheads point to GFP^-^ 4C4^+^ Lcp1^+^ cells. White arrow in (C) and (D) merged channels show erythrocyte nuclei indicating the presence of a vessel. Images were taken using a 20X objective and correspond to orthogonal projections. (n=2). Scale bars 50 µm.

## DISCUSSION

The aim of this study was to perform a detailed characterization of 4C4, a monoclonal antibody that has emerged over the years as a useful tool for immunological investigations in zebrafish and now serves as a gold standard for the prospective detection and/or isolation of microglial cells in this model. Using immunoprecipitation followed by mass spectrometry, we have identified the protein target of 4C4 as Galectin 3 binding protein b, the product of the *lgals3bpb* gene. We have confirmed this identification by assessing antibody binding to HEK-293T cells engineered to express Lgas3bpb. Importantly, our results show that 4C4 is highly specific for Lgals3bpb as it could not recognize recombinant forms of two other Lgals3bp paralogs *in vitro*. Extending previous studies, we also show that 4C4 labels the majority, if not all microglial cells, in both the embryonic and adult brain parenchyma, validating 4C4 as a pan-microglial antibody. Finally, we also provide evidence of 4C4 immunoreactivity outside the brain, including in discrete populations of gut and liver mpeg1^+^ immune cells.

Our identification of Lgals3bpb as a novel marker for zebrafish microglia is supported by several transcriptomic studies, which consistently display strong expression of the gene in bulk populations of embryonic^8^ and adult^18–21^ microglia/macrophages isolated based on fluorescent transgene expression. Recently, *lgals3bpb* transcripts were also found to be highly enriched in microglia in a single-cell transcriptomic profiling of juvenile zebrafish brain immune cells ^8,15,16,20,21^. In comparison, expression of the two paralogous genes *zgc:112492* and *lgals3bpa* in the microglia cluster appeared to be negligible. The microglial expression profile of *lgals3bpb* is also in sharp contrast with that of *Ependymin*, the third protein identified in our IP-MS experiments. The fact that *ependymin* transcripts are barely detected in microglia, together with the low protein sequence conservation with Lgals3bpb, make it an unlikely candidate for 4C4 recognition. Therefore, this protein was excluded from our downstream analyses. As Ependymin is secreted by meningeal fibroblast-like cells in teleost fish^22–24^, its detection in the IP-MS experiment might be explained by its high abundance in the brain extracellular and cerebrospinal fluids ^22–24^.

*Lgals3bpb* is predicted as a zebrafish ortholog of the mammalian LGALS3BP (also known as MAC2-BP or tumor-associated antigen 90K), a member of the family of scavenger receptor cysteine-rich (SRCR) domain-containing proteins, with known intracellular and extracellular functions associated with the immune system. The human and zebrafish protein sequences share approximately 35% identity (not shown) and the structural features of the proteins are conserved in both species (a signal peptide, the SRCR domain, a BTB/POZ (Broad-Complex, Tramtrack and Bric a brac/Poxvirus and Zinc finger) domain, and a BACK (BTB and C-terminal Kelch) domain). In normal conditions, LGALS3BP expression is widely distributed among tissues ^25^ and the protein is also found in serum and body fluids such as saliva or cerebrospinal fluid ^26^.

Our Western-blotting analyses indicate that Lgals3bpb is expressed in a more restricted manner than its mammalian ortholog. Indeed, we found strong 4C4 immuno-reactivity in adult organs containing microglial cells such as brain and eyes, but no signal in an extended panel of tissues. As zebrafish possess two other paralogs, it remains possible that *lgals3bpa* and/or *zgc:112492*, whose distribution patterns are currently unknown, will mark other organs, possibly in a complementary fashion. Nevertheless, despite being predominantly and ubiquitously expressed in microglia, it is clear that *lgals3bpb* is also expressed outside the zebrafish brain^27^, including in macrophages. Therefore, the use of 4C4 to label these cells will depend on the level and extend of *lgals3bpb* expression in the macrophage population in the tissue of interest.

It is known that mammalian microglia can express LGALS3BP. However, while we found zebrafish microglia constitutively express Lgals3bpb in homeostatic conditions, expression of the mammalian ortholog appears mainly to be linked to the disease-associated microglia (DAM) phenotype, a transcriptionally distinct microglial profile shared across various neurodegenerative disorders. For example, LGALS3BP transcripts are upregulated in microglial cells in murine models of Alzheimer’s disease-like, amyotrophic lateral sclerosis (ALS) ^28–30^ and chronic autoimmune diseases ^31^, as well as glioma ^32^ or demyelinated injuries ^33^, suggesting a possible role for LGALS3BP in activated microglia. It will be interesting to investigate whether the divergences of expression observed between species confer different microglia functionalities.

Protein databases (UniProtKB) predict that, similar to its mammalian counterpart ^26,34^, zebrafish Lgals3bpb is highly glycosylated. Interestingly, the protein bands detected by 4C4 in SDS-PAGE have higher apparent molecular weight than that calculated based on Lgals3bpb protein sequence. Given the lack of antibody cross-reactivity with any of the other two Lgals3bpb paralogs, it is tempting to speculate that these protein bands correspond to different Lgals3bpb isoforms resulting from alternative splicing and/or glycosylation. In support of the latter, LGALS3BP calculated molecular weight is approximately 65 kDa and its secreted, glycosylated form is found at 90-100 kDa in Western blotting ^26,34^. Importantly, glycosylation of the mammalian protein is essential for its secretion and interaction with a variety of extracellular signaling molecules (e.g., galectin-1, galectin-3, galectin-7, integrins, collagens, DC-SIGN). Our findings may support a conserved extracellular localization of *lgals3bpb* and a possible role in extracellular matrix interactions between microglia and other cells in the CNS. However, there is also evidence that LGALS3BP possesses intracellular activity, mediated, for example, through interactions with cytoplasmic proteins such as TAK1, a member of the NF-kB pathway ^35,36^. As the roles and modes of action of LGALS3BP in the microglia lineage remain elusive, zebrafish may thus provide a convenient model for the functional dissection of this enigmatic protein in an *in vivo* context.

In summary, our work reveals the identity of the 4C4 protein target and validates this antibody as a useful tool for the prospective identification of microglia during the zebrafish life span. The 4C4 antibody has been previously used for immunofluorescence and flow cytometry ^7,8,37^ and here, we demonstrate its suitability for two additional applications: western blotting and immunoprecipitation. The recognition of both the denatured and native forms of the Lgals3bpb protein by 4C4 indicates that its epitope is linear and accessible in the native protein structure, although additional work will be required to address this question. Identification of Lgals3bpb as the 4C4 antigen will now allow the zebrafish immunology field to move forward on a more solid footing and will open new avenues for understanding the biological functions of this evolutionary conserved microglial marker using the zebrafish model.

## EXPERIMENTAL PROCEDURES

### Zebrafish husbandry

Zebrafish were maintained under standard conditions, according to FELASA ^38^ and institutional (Université Libre de Bruxelles, Brussels, Belgium; ULB) guidelines and regulations. All experimental procedures were approved by the ULB ethical committee for animal welfare (CEBEA) from the ULB. The following transgenic lines were used: Tg(*mpeg1*.*1*:eGFP)^gl22 39^ (here referred to as *mpeg1*:GFP) and TgBAC(*p2ry12*:*p2ry12*-GFP)^hdb3 13^. The csf1r double mutant line used is a combination of *csf1ra*^j4e1 40^ and *csf1rb*^*sa1503*^ mutants (here referred to as *csf1r*^*DM*^) ^41^. Unless specified, the term “adult” fish refers to animals aged between 4 months and 1 year old. For clarity, throughout the text, transgenic animals are referred to without allele designations.

### Hybridoma culture and antibody purification

The hybridoma cell line for 4C4 antibody production was purchased (7.4.C4 ECACC 92092321, Sigma) and maintained in the laboratory following the manufacturer recommended cell culture conditions. For the production and purification of the monoclonal antibody the hybridoma was sent to ProteoGenix (France). The purified version of the antibody was used for the experiments.

### Western blotting

Sample preparation and immunoblotting was performed as previously described ^42^. Briefly, fish were sampled after the experiments and euthanized by immersion in 0.25 mg/ml of buffered MS-222 (Sigma). Individual brains were dissected in PBS and flash frozen in liquid nitrogen and processed immediately or stored at -80°C. Frozen zebrafish adult brains were lysed in RIPA buffer (Sigma), 1mM phenylmethylsulfonyl fluoride (PMSF), 1X protease inhibitor cocktail (Sigma). Protein extracts were quantified using Pierce 660nm Protein Assay (Pierce 22660). 50 µg of total protein was denaturalized in NuPAGE LDS sample buffer (Invitrogen), resolved on a 7.5% SDS-PAGE gel and transferred to a nitrocellulose membrane (Amersham, GE Healthcare). Membranes were blocked, incubated overnight at 4°C with primary antibody, washed and incubated with an anti-rabbit or anti-mouse HRP-conjugated secondary antibody (1:20,000, Invitrogen). Primary antibodies used for western blotting were: 7.4.C4 (1:200) and beta actin (1:1000, Proteintech) was used as a loading control. Immunoreactive bands were developed using enhanced chemiluminescence method (LumiGLO, Cell Signalling) and visualized (Fusion Solo S, FusionCapt Advance Solo software).

### Immunoprecipitation

Frozen zebrafish adult brains (n = 50) were lysed in NP-40 buffer (25 mM Tris HCl ph7.5, NaCl 75 mM, NP-40 0.5%, NaF 25 mM) with 10% glycerol, 1 mM DTT and protease inhibitors (1 mM PMSF, 1X protease inhibitor cocktail). Lysates were quantified as described in the previous section. Thirty microliters of protein G/protein A agarose beads (EMD Millipore) were washed with lysis buffer (containing protein inhibitors) and incubated with 5 µg of 4C4 antibody or IgG1^k^ as a control isotype (ab18443 Abcam) at 4°C overnight under rotation. Next, agarose beads were washed and incubated with 5 mg of brain protein lysate during 5h at 4°C under rotation and washed with lysis buffer. For WB analyses, agarose beads were resuspended in 100 µl of NuPAGE LDS sample buffer (Invitrogen). For MS analysis, agarose beads were washed in lysis buffer without NP-40, washed once in milliq water, dried, flash frozen in liquid nitrogen, and stored at -80°C until processing.

### Liquid chromatography-Mass Spectrometry (LC-MS/MS)

Dried agarose beads were resuspended in SDC buffer (sodium deoxycholate 1%, 10 mM TCEP, 55 mM chloroacetamide, 100 mM Tris HCl pH8.5) and denatured during 10 minutes at 95°C. Samples were diluted two-fold with 100 mM of triethylammonium bicarbonate (TEAB) and proteins were digested during 3 hours with 1 µg of trypsin (Promega V5111) and 1 µg of LysC (Wako 129-02541) at 37°C. Peptides were purified using SDB-RPS columns (Affinisep). Briefly, digested peptides were diluted two-fold with 2% TFA/isopropanol, mixed thoroughly and loaded on a SDB-RPS column. After washing (1% TFA/isopropanol followed by 5% ACN/0.2% TFA), peptides were eluted with 5% NH4OH/60% ACN and evaporated to dryness at 45°C. 80% of resuspended peptides (8/10 µl in 100% H2O/0.1% HCOOH) were injected on a Triple TOF 5600 mass spectrometer (Sciex, Concord, Canada) interfaced to an EK425 HPLC System (Eksigent, Dublin, CA) and data were acquired using Data-Dependent-Acquisition (DDA). Peptides were injected on a separation column (Eksigent ChromXP C18, 150 mm, 3 µm, 120 A) using a two steps acetonitrile gradient (5-25% ACN/0.1% HCOOH in 48 min then 25%-60% ACN/0.1% HCOOH in 20 min at 5 µl/min) and were sprayed online in the mass spectrometer. MS1 spectra were collected in the range 400-1250 m/z with an accumulation of 250 ms. The 20 most intense precursors with a charge state 2-4 were selected for fragmentation, and MS2 spectra were collected in the range of 50-2000 m/z with an accumulation of 100 ms; precursor ions were excluded for reselection for 12 s.

### MS data analysis

Raw data were analyzed using Fragpipe computational platform (v15.0) with MSfragger ^43^ (v3.2), Philosopher ^44^ (v3.4.13; build 1611589727) and IonQuant ^45^ Peptides identifications were obtained using MSFragger search engine on .mzML files, from converted .Wiff/Wiff.scan files, on a protein sequence database of zebrafish (UP0000004372021) from Uniprot (downloaded 24th Feb, 2021, containing “sp” and “tr” sequences, no isoforms) supplemented with common contaminant proteins and reversed protein sequences as decoys. Mass tolerances for precursors and fragments were set to 30 and 20 ppm respectively, and with spectrum deisotoping mass calibration ^46^, and parameter optimization enabled. Enzyme specificity was set to “trypsin” with enzymatic cleavage and a maximum of 5 missed trypsin cleavages were allowed. Isotope error was set to 0/1/2. Peptide length was set from 6 to 50, and peptide mass was set from 500 to 5000 Da. Variable modifications (methionine oxidation, acetylation of protein N-termini, and pyro-Glu [-17.0265 Da]) were added while carbamidomethylation of Cysteine was set as a fixed modification. Maximum number of variable modifications per peptide was set to 5. MS/MS search results were further processed using the Philosopher toolkit with PeptideProphet (with options for accurate mass model binning, semi-parametric modeling with computation of possible non-zero probabilities for decoy entries) and with ProteinProphet. Further filtering to 1% protein-level FDR allowing unique and razor peptides were used and final generated reports were filtered at each level (PSM, ion, peptide, and protein) at 1% FDR. Label free quantification was performed using IonQuant with MBR and normalization enabled, 2 ions minimal and default options. Further statistical analysis and visualization were performed in R (v4.1) using commonly used packages. Distribution of protein intensity were normalized using a quantile normalization method (apmsWapp::norm.inttable v1.0) and enrichment was calculated for each pair of IP (4C4 vs IgG CTL), averaged and subjected to a paired t-test.

Multiple sequence alignment of the sequence of the 3 top hits and of its paralogs was generated using Clustal Omega webtool (www.ebi.ac.uk/Tools/msa/clustalo/) ^47^, visualized using Jalview and manually annotated.

### Cloning and transient expression of *lgals3bpb* and zgc:*112492*

The 1719 bp coding sequence (CDS) of the *lgals3bpb* (ENSDARG00000040528), was amplified using a high-proof reading polymerase (CloneAmp HiFi, Takara) with primers containing *EcoRI* and *NotI* restriction sites and subcloned into pCR blunt II TOPO vector for subsequent restriction and ligation into the pcDNA3 expression vector. The cloned CDS was compared with the original and no amino acid mutations were found. The 1686 bp CDS of the zgc:*112492* was synthetized and cloned into pcDNA3 containing *KpnI* and *EcoRI* restrictions sites (GeneCust, France). The human embryonic kidney (HEK293T) cells were cultured in Dulbecco’s modified Eagle’s medium (DMEM) supplemented with 10% FBS and transfected with Lipofectamine 2000 (Invitrogen) according to manufacturer’s instructions. For immunofluorescence, 100 ng of pcDNA3 empty or pcDNA3-*lgals3bpb*, pcDNA3-*zgc:112492*, were used to transfect 1.25×10^5^ cells plated on 0.1% gelatin-coated coverslips in 24-well plates. HEK293T cells were fixed 2 days after transfection in 4% PFA for 30min at RT, washed three times in PBS and stored at 4°C to perform immunostaining (see below).

### Immunostaining and imaging

Adult tissues were dissected, fixed in 4% PFA, incubated overnight in 30% sucrose:PBS before snap-freezing in OCT (Tissue-Tek, Leica) and stored at -80°C. Immunostaining was performed on 14 µm cryosections as described ^11^. For HEK293 cells, a blocking step of 30 min at RT was performed (3% BSA, 5% donkey/sheep serum, 0.3% Triton X-100) before incubation with the mouse 4C4 monoclonal antibody overnight at 4°C. Cells were washed 3 times in PBS and incubated with a mouse secondary antibody (1:500, Abcam) and DAPI (1:1000, Thermofisher). The following primary and secondary antibodies were used: chicken anti-GFP polyclonal antibody (1:500; Abcam), rabbit anti-DsRed polyclonal antibody (1:500; Clontech), rabbit anti-Lcp1 (1:1000), mouse 4C4 monoclonal antibody (1:200), Alexa Fluor 488-conjugated anti-chicken IgG antibody (1:500; Invitrogen), Alexa Fluor 488-conjugated anti-mouse IgG antibody (1:500; Invitrogen), Alexa Fluor 594-conjugated anti-rabbit IgG (1:500; Abcam) and Alexa Fluor 647-conjugated anti-mouse IgG (1:500; Abcam).

Imaging was performed on a confocal Zeiss LSM 780 inverted microscope, using a Plan Apochromat 20× objective for adult sections and a LDLCI Plan Apochromat 25× water-immersion objective for whole-mount embryos and tissue-cleared brains. For HEK293 cells, confocal images were acquired using a Plan Apochromat 20x or LDC Apochromat 40x objective using numerical zoom, as indicated in the figure legends.

## ACKNOWLEDGEMENTS

This study was funded by the Fonds de la Recherche Scientifique (FNRS) (F451218F, UN06119F and UG03019F to V.W., and 1236220F to M.R.) and the Minerve Foundation (to V.W.). This work was also supported by a Research Fellowship from the FNRS (to A.M.), and fellowships from the Erasme Fund (to L.C.), the Belgian Kid’s Fund (to M.M.), and the Televie (to J.P.). This project has also received funding from the European Union’s Horizon 2020 research and innovation programme under the Marie Skłodowska-Curie grant agreement No 843107 (to X.B.)and from the Région de Bruxelles Capitale – Innoviris (RBC/BFB 1 to X.B.) We thank Marianne Caron for technical assistance and members of the Wittamer lab for critical discussion and comments on the manuscript.

## SUPPLEMENTAL MATERIAL

Supplementary Table S1 (xlsx): MS results.

